# DeepATAC: A deep-learning method to predict regulatory factor binding activity from ATAC-seq signals

**DOI:** 10.1101/172767

**Authors:** Naozumi Hiranuma, Scott Lundberg, Su-In Lee

## Abstract

Determining the binding locations of *regulatory factors*, such as transcription factors and histone modifications, is essential to both basic biology research and many clinical applications. Obtaining such genome-wide location maps directly is often invasive and resource-intensive, so it is common to impute binding locations from DNA sequence or measures of chromatin accessibility. We introduce DeepATAC, a deep-learning approach for imputing binding locations that uses both DNA sequence and chromatin accessibility as measured by ATAC-seq. DeepATAC significantly outperforms current approaches such as FIMO motif predictions overlapped with ATAC-seq peaks, and models based only on DNA sequence, such as DeepSEA. Visualizing the input importances for the DeepATAC model reveals DNA sequence motifs and ATAC-seq signal patterns that are important for predicting binding events. The Keras implementation and analysis pipelines of DeepATAC are available at https://github.com/hiranumn/deepatac.

## 1 Introduction

Knowing when and where proteins such as transcription factors interact with DNA is important for both clinical and research purposes. Biological assays such as ChIP-seq [5] are designed to directly measure these interactions, but they require significant resources and a large biological sample. To address these limitations several methods have been proposed to impute these binding locations from raw DNA sequence. The FIMO software [4] from the MEME suite, and deep-learning approaches such as DeepSea [11] and Basset [6] have been successful in this regard. However, DNA sequence alone does not contain any cell-type specific information, which is important for making more accurate predictions. This motivates combining DNA based predictions with predictions from cell type specific data sources.

There are several biological assays that measure cell type specific organization of genome, namely, DNase-seq, MNase-seq, HiC, and ATAC-seq [2, 7, 1, 3]. ATAC-seq is the most recent method and is rapidly gaining popularity due to its cost-efficiency and simplicity. In particular, ATAC-seq only requires 500 to 50,000 cells to measure in-vivo open chromatin signal, while other assays require millions of cells. This is a particularly important feature in clinical settings where you cannot sample a large number of cells from patients when performing personal level analysis. Traditionally, researchers have used putative binding locations predicted by motif finding algorithms (e.g FIMO) overlapped with ATAC-seq peaks to determine where transcription factors are bound. DeepATAC, a deep-learning model that is jointly trained on both ATAC-seq and DNA sequence, significantly outperforms this traditional approach.

## 2 Results

To evaluate DeepATAC, we compared with FIMO [4] predictions overlapped with ATAC-seq peaks, and with DeepSEA [11] DNA based predictions. We use subsets of ChIP-seq binding signals obtained from the DeepSEA project, which originally came from ENCODE database, as true labels for evaluation. We then calculate area under precision recall curves (AUPRC) for 163 and 91 regulatory factors for K562 and GM12878 cell types, respectively.

Figure 1a shows the performance of the three models on predicting CTCF, GATA2, and STAT1 binding events in K562. For all three factors, DeepATAC significantly outperformed the two baseline methods. For STAT1, DeepSEA was unable to learn sequence motifs of the transcription factor and under-performed when compared to FIMO predictions overlapped with ATACseq peaks. On the other hand, DeepATAC managed to predict the binding sites more accurately with the help of the ATAC-seq signals.

**Figure 1:**
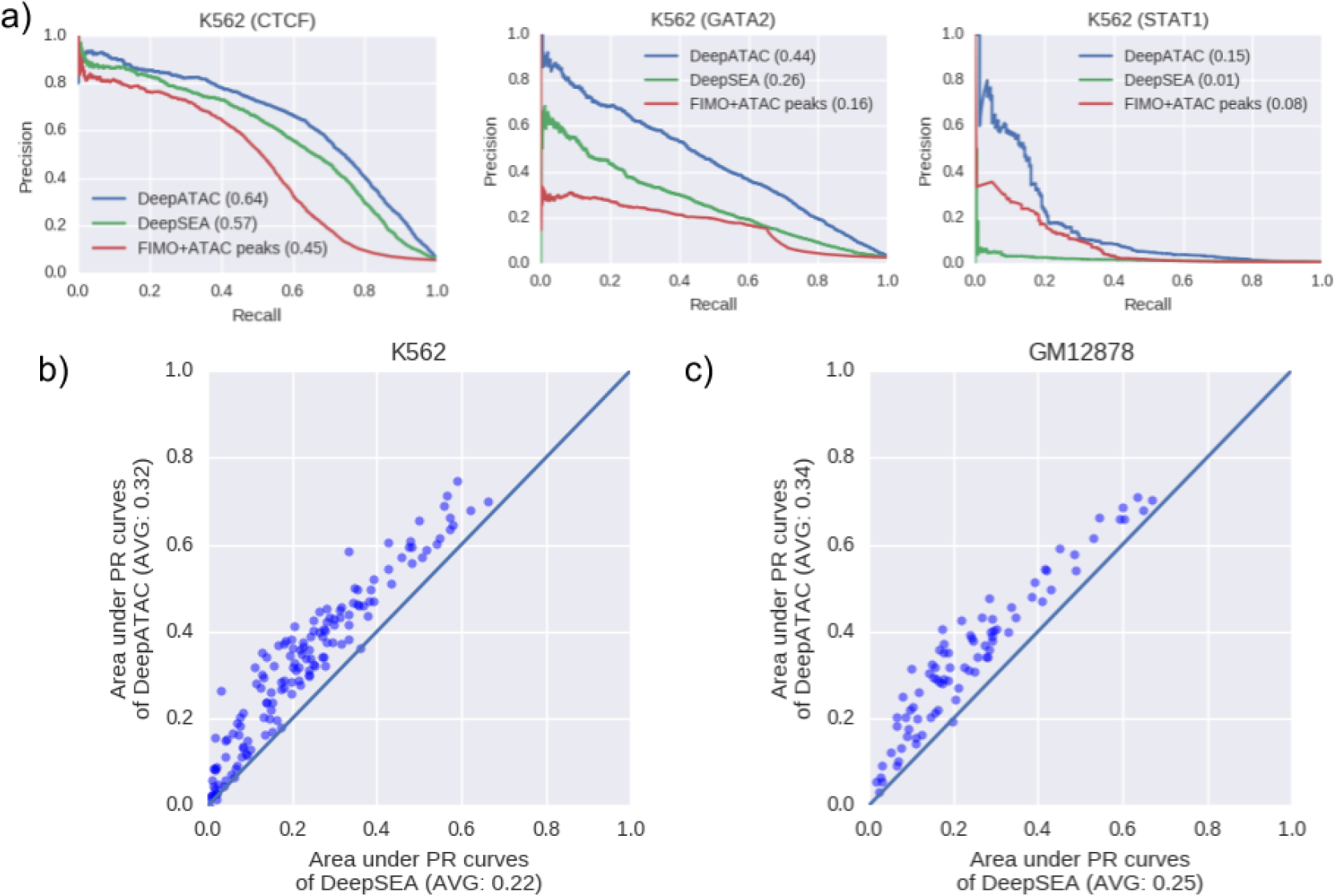
a) Precision recall curves for three transcription factors (CTCF, GATA2, and STAT1) in K562. Areas under the curves (AUPRCs) are shown in parentheses, and the FIMO prediction is masked by ATAC-seq peaks. b-c) A scatter plot comparing the AUPRCs of DeepATAC and DeepSEA in K562 (b) and GM12878 (c). Each data point represents a transcription factor, and points fall above the diagonal when DeepATAC outperforms DeepSEA.

DeepATAC outperformed DeepSEA in 159 out of 163 factors (98%) in K562 with a mean difference in area under PR curves of 0.1 (Figure 1b). Similarly, in GM12878, DeepATAC surpassed DeepSEA in 89 out of 91 factors (97%) with a mean difference of 0.09 (Figure 1c). This represents more than a 40% improvement over DeepSEA predictions. DeepATAC continued to outperform DeepSEA even when the samples were restricted to only fall within ATAC-seq peak regions (146 out of 163 in K562 and 84 out of 91 in GM12878), indicating that DeepATAC capture subtle ATAC-seq signals lost during peak calling. DeepATAC also outperformed the FIMO predictions within ATAC-seq peak regions in 38 out of 39 factors in K562 and 18 out of 20 factors in GM12878.

To investigate what signals DeepATAC was using to make predictions, we used integrated gradients, a feature attribution method, to determine which sequence and ATAC-seq signal patterns are informative for predicting transcription factor binding sites [9]. Figure 2 shows that Deep-ATAC was able to learn both forward and reverse CTCF motifs to make the correct predictions. The attribution values for ATAC-seq signals indicate that the presence of 5-end cut sites outside of the learned motifs have positive contributions to the correct prediction, and the absence of the cut sites within the motifs have negative contributions, suggesting the the model may be able to capture signals as subtle as ATAC-seq footprints.

**Figure 2:**
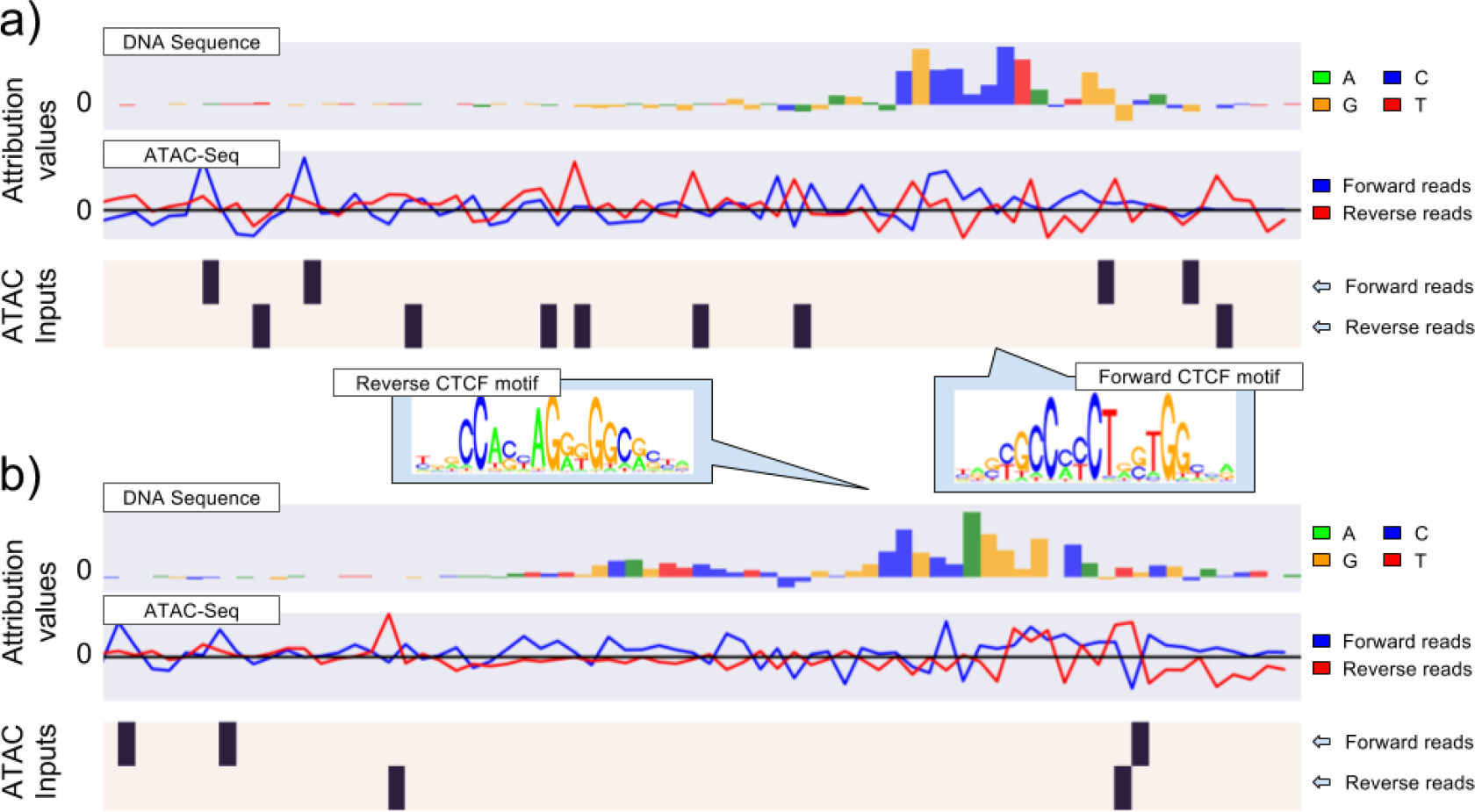
Attribution values generated using integrated gradients for true positive CTCF locations in K562 predicted by DeepATAC with high confidence (probability *>* 0.95). a-b) The first row is a bar plot of attribution values for the DNA sequence, where green, yellow, blue, and red bars correspond to A, G, C, and T, respectively. The second row is the attribution values for the presence of ATAC-seq 5’-end cut sites for forward and reverse reads. The third row is the binarized ATAC-seq inputs for these samples. The integrated gradients reference was 0 for DNA inputs and 0.5 for ATAC-seq.

## 3 Discussion

DeepATAC is a powerful tool to predict the location of genome-wide regulatory activity. It outperforms common approaches such as FIMO or DeepSEA by successfully integrating DNA sequence and ATAC-seq signal. This integration provides a 40% average performance improvement over DeepSEA in K562 and GM12878. It appears that DeepATAC not only captures the enrichment of binding activity in ATAC-seq signals, but also captures subtle signals such as footprints to make more accurate predictions. Looking forward, we hope to extend our work to cross-celltype predictions. If we can learn complex and generalizable interactions between ATAC-seq and DNA sequence across many different cell types, we may be able to apply the model to make personalized predictions for disease patients across many cell types, since SNP information and ATAC-seq signals are becoming increasingly easier to obtain. One can then use the predicted regulatory activities to infer information such as drug sensitivity.

## 4 Methods

### Data

The DeepATAC model was trained on the subset of DeepSEA data [11]. Specifically, the DNA sequence samples were 1 hot-encoded 200 base pair windows (4 by 200 matrix), where we observed (using ChIP-seq peaks) at least one of the 163 and 91 regulatory factors for the K562 and GM12878 cell types respectively. ATAC-seq data was obtained from Greenleaf et al [3] and Schimidt et al [8]. For each cell type, we pooled all ATAC-seq reads and removed duplicates. We then used Bowtie2 to map reads to the hg19 genome. For each sample, we generated a 2 by 100 matrix containing 5-end cut sites of reverse and forward reads. Concatenated with the DNA sequence sample, this forms a 6 by 100 matrix per sample. The test samples were generated from chromosome 8 and 9, and the training and validation samples were generated from other chromosomes except for X and Y.

### Deep learning architecture

Both DeepATAC and DeepSEA models consist of three convolution layers and two dense layers [11] (Figure 3). Each convolution is implemented by stacking a convolution layer with a kernel size of 8 base pairs, a batch normalization layer, a rectified linear unit (Relu), a max pool layer, and a drop out layer. These three convolution layers have 160, 240, and 480 kernels, respectively. The dropout probabilities are set to 0.2, 0.2, and 0.5. In addition, minor L2 regularization (5e-07) is applied to convolution kernels. The first dense layer has 600 hidden units with Relu activations, and the second dense layer has 163 and 91 nodes (K562 and GM12878 respectively) with sigmoid activations. Minor L1 regularization of 1e-08 is added to the dense weights. The training was performed with an Adam optimizer for 30 epochs. The best models were chosen based on validation performance. Keras models are available at https://github.com/hiranumn/deepatac.

**Figure 3:**
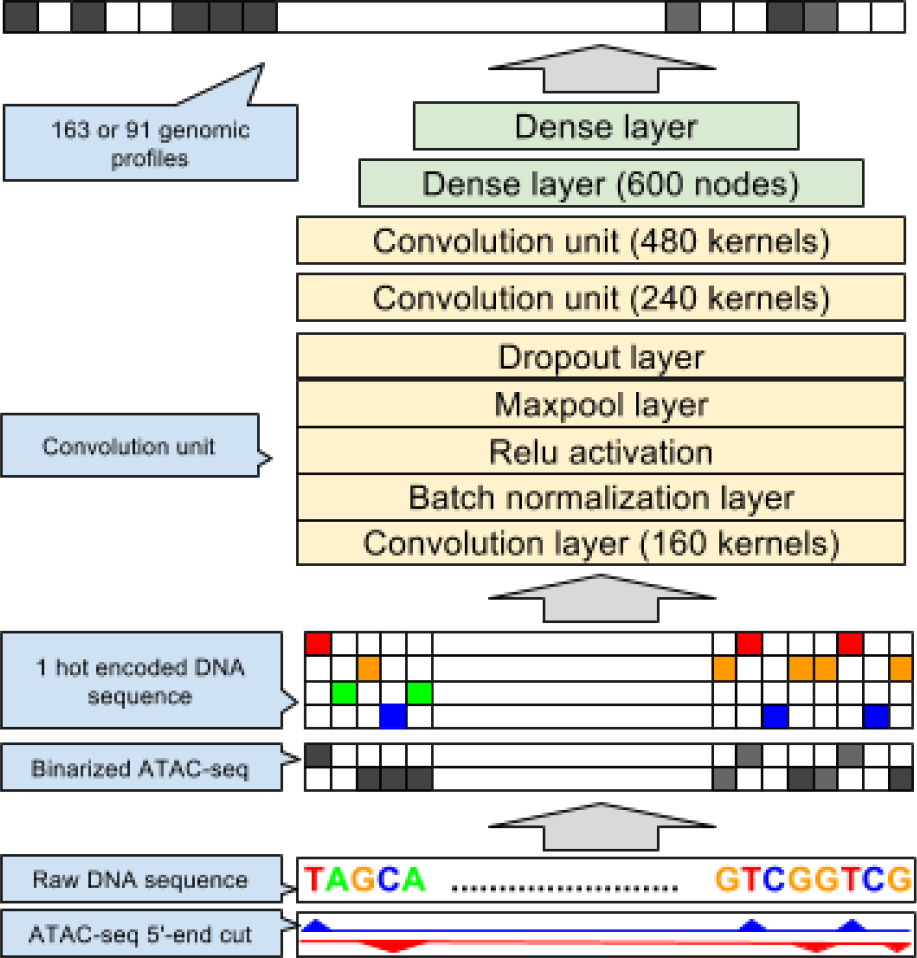
The deep learning architecture for DeepATAC.

### Other materials

FIMO predictions were obtained by running FIMO with a q-value cutoff of 0.05 on JASPAR vertebrate motifs downloaded from the MEME suite website at http://memesuite.org/db/motifs. ATAC-seq peaks were called by MACS2 [10] with the “–*broad*” option and a p-value cutoff of 0.01. We used an integrated gradient framework [9] to interpret and visualize our model. Our python implementation of integrated gradients for Keras can be found at https://github.com/hiranumn/IntegratedGradients.

